# Reconstitution of cargo-induced LC3 lipidation in mammalian selective autophagy

**DOI:** 10.1101/2021.01.08.425958

**Authors:** Chunmei Chang, Xiaoshan Shi, Liv E. Jensen, Adam L. Yokom, Dorotea Fracchiolla, Sascha Martens, James H. Hurley

## Abstract

Selective autophagy of damaged mitochondria, intracellular pathogens, protein aggregates, endoplasmic reticulum, and other large cargoes is essential for health. The presence of cargo initiates phagophore biogenesis, which entails the conjugation of ATG8/LC3 family proteins to membrane phosphatidylethanolamine. Current models suggest that the presence of clustered ubiquitin chains on a cargo triggers a cascade of interactions from autophagic cargo receptors through the autophagy core complexes ULK1 and class III PI 3-kinase complex I (PI3KC3-C1), WIPI2, and the ATG7, ATG3, and ATG12-ATG5-ATG16L1 machinery of LC3 lipidation. This model was tested using giant unilamellar vesicles (GUVs), GST-Ub_4_ as a model cargo, the cargo receptors NDP52, TAX1BP1, and OPTN, and the autophagy core complexes. All three cargo receptors potently stimulated LC3 lipidation on GUVs. NDP52- and TAX1BP1-induced LC3 lipidation required the ULK1 complex together with all other components, however, ULK1 kinase activity was dispensable. In contrast, OPTN bypassed the ULK1 requirement completely. These data show that the cargo-dependent stimulation of LC3 lipidation is a common property of multiple autophagic cargo receptors, yet the details of core complex engagement vary considerably and unexpectedly between the different receptors.

## Introduction

Macroautophagy (hereafter autophagy) is an evolutionarily conserved catabolic pathway that sequesters intracellular components in double membrane vesicles, autophagosomes, and delivers them to lysosomes for degradation (*1*). By removing excess or harmful materials like damaged mitochondria, protein aggregates, and invading pathogens, autophagy maintains cellular homeostasis and is cytoprotective (*2*). Autophagy is particularly important in maintaining the health of neurons, which are long-lived cells that have a high flux of membrane traffic. Defective autophagy of mitochondria (mitophagy) downstream of mutations in PINK1 and Parkin is thought to contribute to the etiology of a subset of Parkinson’s Disease (*3*).

The *de novo* formation of autophagosome, central to autophagy, entails the formation of a membrane precursor, termed the phagophore (or isolation membrane) that expands and seals around cytosolic cargoes (*4*). A set of autophagy related (ATG) proteins drive autophagosome biogenesis. In mammalian cells, the unc-51-like kinase 1 (ULK1) complex, consisting of ULK1 itself, FIP200, ATG13, and ATG101, is typically recruited to the autophagosome formation site first. The class III phosphatidylinositol 3-kinase complex I (PI3KC3-C1) is subsequently activated to generate phosphatidylinositol-3-phosphate (PI(3)P). PI(3)P enriched membranes serve as platforms to recruit the downstream effector WIPIs (WD-repeat protein interacting with phosphoinositides), and ATG8/LC3 conjugation machinery (*5, 6*). ATG2 transfers phospholipids from endoplasmic reticulum (ER) to the growing phagophore (*7–9*), while ATG9 translocates phospholipids from the cytoplasmic to the luminal leaflet, enabling phagophore expansion (*10, 11*). These above-mentioned proteins are sometimes referred to as the “core complexes” of autophagy.

The attachment of the ATG8 proteins of the LC3 and GABARAP subfamilies to the membrane lipid phosphatidylethanolamine (PE), termed LC3 lipidation, is a hallmark of autophagosome biogenesis. LC3 lipidation occurs via a ubiquitin-like conjugation cascade. The ubiquitin E1-like ATG7 and the E2-like ATG3 carry out the cognate reactions in the LC3 pathway. The ATG12-ATG5-ATG16L1 complex scaffolds transfer of LC3 from ATG3 to PE (*12, 13*). The role of the ATG12-ATG5-ATG16L1 is analogous to that of a RING domain ubiquitin E3 ligase, although there is no sequence homology between any of the subunits and ubiquitin E3 ligases. Covalent anchoring of LC3 to membrane is closely associated with phagophore membrane expansion (*14–17*) and cargo sequestration (*18–20*). Although recent evidence showed that autophagosome formation can still occur in mammalian cells lacking all six ATG8 family proteins (*17, 21*), their size was reduced and lysosomal fusion was impaired. LC3 lipidation is thus involved in multiple steps in autophagosome biogenesis, and is critical for promoting autophagosome-lysosome fusion (*17, 21*).

In most instances in mammalian cells, autophagy is highly selective and tightly regulated (*22*). Several targets of selective autophagy have been described, including mitochondria (mitophagy), intracellular pathogens (xenophagy), aggregated proteins (aggrephagy), endoplasmic reticulum (reticulophagy), lipid droplets (lipophagy), and peroxisomes (pexophagy) (*23*). The achievement of selectivity relies on a family of autophagy receptors, which specifically bind to cargoes and the phagophore (*24–26*). Some types of selective autophagy like aggrephagy, mitophagy, and xenophagy are initiated by the ubiquitination of cargoes, which are recognized by a subset of cargo receptors including p62 (sequestosome-1), NBR1, optineurin (OPTN), NDP52 and Tax1-binding protein 1 (TAX1BP1). All of these receptors contain a LC3-interaction region (LIR), a ubiquitin binding domain (UBD), and a dimerization/oligomerization domain (*18, 26, 27*). These cargo receptors are well-known to connect cargo to the phagophore through their interaction with both clustered ubiquitin chains and membrane-conjugated LC3. Several cargo receptors have recently been shown to trigger autophagy initiation, thus functioning upstream of LC3 lipidation. NDP52 directly binds to and recruits the ULK1 complex to damaged mitochondria and intracellular bacteria by binding to the coiled-coil (CC) of the FIP200 subunit of the ULK1 complex (*28–30*). p62 is also recruited to FIP200, but to its CLAW domain instead of the CC (*31*). Initiation of mitophagy by OPTN, however, appears to be independent of ULK1 (*32*). These findings are beginning to reveal different roles for various cargo receptors in triggering early autophagy machinery assembly via distinct entry points.

*In vitro* reconstitution studies have recently shown to recapitulate the steps in autophagosome formation, especially in yeast autophagy (*33–36*). These *in vitro* approaches are powerful to investigate the molecular mechanisms of such a complicated cell biological process by controlling the multi-component compositions and spatiotemporal arrangements. However, it has been challenging to reconstitute mammalian autophagosome because of the complexity of mammalian autophagy machinery. As part of a long-term effort, we recently reconstituted the events from PI(3)P production by the PI3KC3-C1 to LC3 lipidation in mammalian autophagy in a giant unilamellar vesicle (GUV) model system (*37*). Separately, we showed that in the presence of clustered ubiquitin chains, NDP52 promotes recruitment of the ULK1 complex to membranes (*29*). Here, using the GUV model system, and focusing on autophagy receptors involved in mitophagy, we established a start-to-finish reconstitution of selective autophagy initiation from autophagy receptor engagement through LC3 lipidation. We found that NDP52, TAX1BP1 and OPTN triggered robust LC3 lipidation in the presence of the ULK1 complex, PI3KC3-C1 complex, WIPI2, and LC3 conjugation machinery. LC3 lipidation triggered by NDP52 and TAX1BP1 was dependent on both ULK1 and PI3KC3-C1, while OPTN-induced LC3 lipidation was only dependent on the activity of PI3KC3-C1. We further found that these cargo receptors trigger LC3 lipidation through distinct multivalent webs of interactions, thereby enabling the rapid LC3 lipidation for autophagosome formation.

## Results

### Reconstitution of NDP52 and TAX1BP1-triggered LC3 lipidation

We sought to establish a purified system that recapitulate the initiation of mitophagy, which is known to utilize the cargo receptors NDP52, TAX1BP1, and OPTN (*38–43*), together with the core autophagy initiation machinery and intracellular membranes. We used GUVs with an ER-like lipid composition to mimic the membranes, a mixture of linear tetraubiquitin and cargo receptors to mimic cargo signals, and these were incubated with a set of purified core autophagy machineries that are involved in autophagy initiation, including the ULK1 complex, the PI3KC3-C1 complex, the PI(3)P effector WIPI2d, the E1-related ATG7, the E2-related ATG3, the functional ubiquitin E3 ligase counterpart ATG12-ATG5-ATG16L1 complex (hereafter referred to as “E3” or “E3 complex” for brevity), and LC3B (Fig. 1A and fig. S1). All proteins and complexes used were full-length and wild-type, with the exception of the FIP200 Δ641-779 construct (fig. S2A), which was engineered to increase stability (fig. S2B) and prevent non-specific aggregation. Negative stain electron microscopy (NSEM) images showed that FIP200^Δ641-779^ had essentially the same structure as wild-type, while losing its propensity to aggregate (fig. S2C) (*29*). Contour lengths and end-to-end distances of FIP200^Δ641-779^ as analyzed by NSEM were comparable as the full-length (fig. S2D) (*29*). All of the fluorescently tagged fusion constructs were previously characterized and shown to be functional (*29, 37*). The typical concentrations of autophagy proteins in human cells are unknown, but as most are thought to be scarce, we used the following concentrations for all reconstitution reactions: 5 μM GST-Ub_4_, 500 nM cargo receptors, 25 nM ULK1 complex, 25 nM PI3KC3-C1 complex, 100 nM WIPI2d, 50 nM E3 complex, 100 nM ATG7, 100 nM ATG3, and 500 nM mCherry-LC3B.

**Fig.1.**
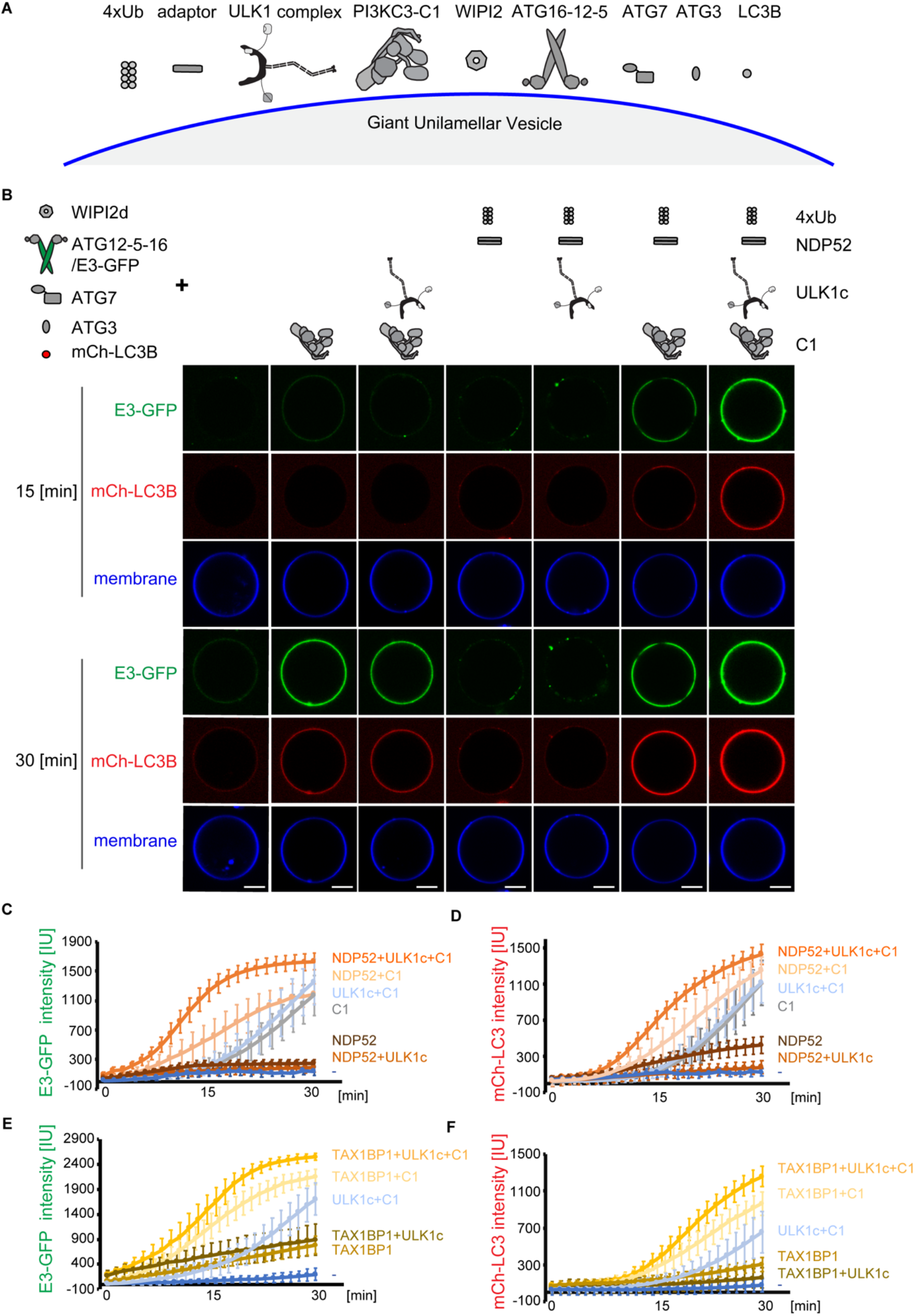
Reconstitution of NDP52 and TAX1BP1-triggered LC3 lipidation. (A) The schematic drawing illustrates the reaction setting. The blue curve indicates the GUV membrane. Gray cartoons are autophagy components present in the reaction. (B) Representative confocal images showing the membrane recruitment of E3 complex (green) and LC3B (red). GUVs were incubated with WIPI2d, E3-GFP, ATG7, ATG3, mCherry-LC3B, ATP/Mn^2+^, and different upstream components as listed above each image column. Images taken at 15 min and 30 min are shown. Scale bars, 10 μm. (C and D) Quantitation of the kinetics of mCherry-LC3B (C) and E3-GFP (D) recruitment to the membrane from individual GUV tracing in A (Averages of 50 vesicles are shown, error bars indicate standard deviations). (E and F) GUVs were incubated with WIPI2d, E3-GFP, ATG7, ATG3, mCherry-LC3B, ATP/Mn^2+^, and different proteins listed above the images in Fig.S3. Quantitation of the kinetics of mCherry-LC3B (E) and E3-GFP (F) recruitment to the membrane from individual GUV tracing (Averages of 50 vesicles are shown, error bars indicate standard deviations). All results representative of three independent experiments.

We first investigated the recruitment of the E3 complex to membranes, since the localization of E3 complex dictates the sites of LC3 lipidation in cells (*44*). We observed that in the presence of WIPI2d and the LC3 conjugation machinery, but not ULK1 complex or PI3KC3-C1, little or no GFP-E3 or mCherry-LC3B was recruited to the GUVs within 30 min (Fig. 1B, first column). The addition of PI3KC3-C1 triggered the membrane recruitment of both E3 and LC3B (Fig. 1B, second column), consistent with the previous observation that PI3KC3-C1 activity is required for E3 membrane targeting and LC3 lipidation (*37*). ULK1 phosphorylates core subunits of PI3KC3-C1 (*45, 46*), but the addition of ULK1 complex together with PI3KC3-C1 had similar effects to PI3KC3-C1 alone (Fig. 1B, third column). Because NDP52 and the ULK1 complex have been shown to interact directly with the E3 complex (*47–49*), we asked whether NDP52 or ULK1 could mediate the membrane recruitment of E3 complex. However, the addition of NDP52 with GST-Ub_4_ or that together with ULK1 complex did not result in an obvious increase of membrane enrichment of E3 and subsequent LC3B (Fig. 1B, fourth and fifth columns). However, NDP52 and GST-Ub_4_ did enhance PI3KC3-C1 triggered E3 and LC3B membrane recruitment (Fig. 1B, sixth column). Membrane recruitment was further enhanced when ULK1 complex was added (Fig. 1B, last column).

Quantification of the kinetics of membrane binding showed that both E3 recruitment and LC3 lipidation were fastest when all the components were present (Fig. 1C and D). The kinetics of E3 recruitment to membranes were faster than that of LC3 (Fig. 11C and D), indicating that the E3 is being recruited in a catalytically competent form. We found that LC3 lipidation was slightly more efficient in the presence of the FIP200^Δ641-779^ version of the ULK1 complex relative to the version containing wild-type FIP200 (fig. S2E). We thus used the FIP200^Δ641-779^ ULK1 complex for all further reconstitution assays and refer to it hereafter as simply the “ULK1 complex”. Taken together, these data show that NDP52 triggers efficient LC3 lipidation when both ULK1 and PI3KC3-C1 complexes are present.

We next tested TAX1BP1, a structural paralog of NDP52 which has roles in mitophagy, xenophagy, and aggrephagy (*42, 50, 51*). We found that similar to NDP52, TAX1BP1 induced the most robust and efficient E3 membrane binding and LC3 lipidation when both ULK1 and PI3KC3-C1 complex were included in the system (Fig. 1E and F and fig. S3). Although the addition of TAX1BP1 and GST-Ub_4_ resulted in a slightly stronger E3 recruitment than NDP52, no obvious increase in LC3 lipidation was observed. (Fig. 1F and fig. S3, fourth and fifth columns). Together, these data indicate that, like NDP52, TAX1BP1 can trigger robust LC3 lipidation in response to a cargo mimetic *in vitro*, which is dependent on both ULK1 and PI3KC3-C1 complex.

### Reconstitution of OPTN triggered LC3 lipidation

We went on to investigate another cargo receptor OPTN, which has been shown to mediate Parkin-dependent mitophagy (*39*). Residues S177 and S473 of OPTN are phosphorylated by Tank-binding kinase 1 (TBK1), which were reported to enhance the binding of OPTN to both LC3 and ubiquitin (*52*). We first tested the phosphomimetic double mutant of OPTN S177D/S473D, hereafter “OPTNS2D”. We observed that OPTNS2D and GST-Ub_4_ alone induced a modest recruitment of both E3 and LC3 to the GUV membrane, similar to the addition of PI3KC3-C1 (Fig. 2A, first four columns). In addition, OPTNS2D and GST-Ub_4_ dramatically increased E3 recruitment and LC3 lipidation by PI3KC3-C1 (Fig. 2A, sixth column). However, in contrast to the situation with NDP52 or TAX1BP1, the addition of ULK1 complex had no effect on either E3 or LC3 binding triggered by the OPTNS2D-Ub-PI3KC3-C1 axis (Fig. 2A, last column). The dynamics of LC3 lipidation and E3 binding when OPTNS2D, GST-Ub_4_ and PI3KC3-C1 were present were essentially the same in the presence or absence of ULK1 complex (Fig. 2B and C). These data indicate that OPTNS2D can also trigger a robust LC3 lipidation, but as distinct from NDP52 and TAX1BP1, OPTN triggered LC3 lipidation depended only on the activity of PI3KC3-C1. We compared the kinetics of E3 recruitment compared to that of LC3 in the presence of all components, LC3 was recruited slower than E3 with a mean lag of 4.5 min (fig. S4). We also evaluated the activity of wild type OPTN in the presence or absence of PI3KC3-C1. OPTNWT and GST-Ub_4_ also enhanced membrane binding of E3 and LC3 lipidation, but more weakly than OPTNS2D (Fig. 2D and E), indicating that the higher affinities for LC3 and ubiquitin contributed to faster LC3 lipidation in the presence of OPTNS2D.

**Fig. 2.**
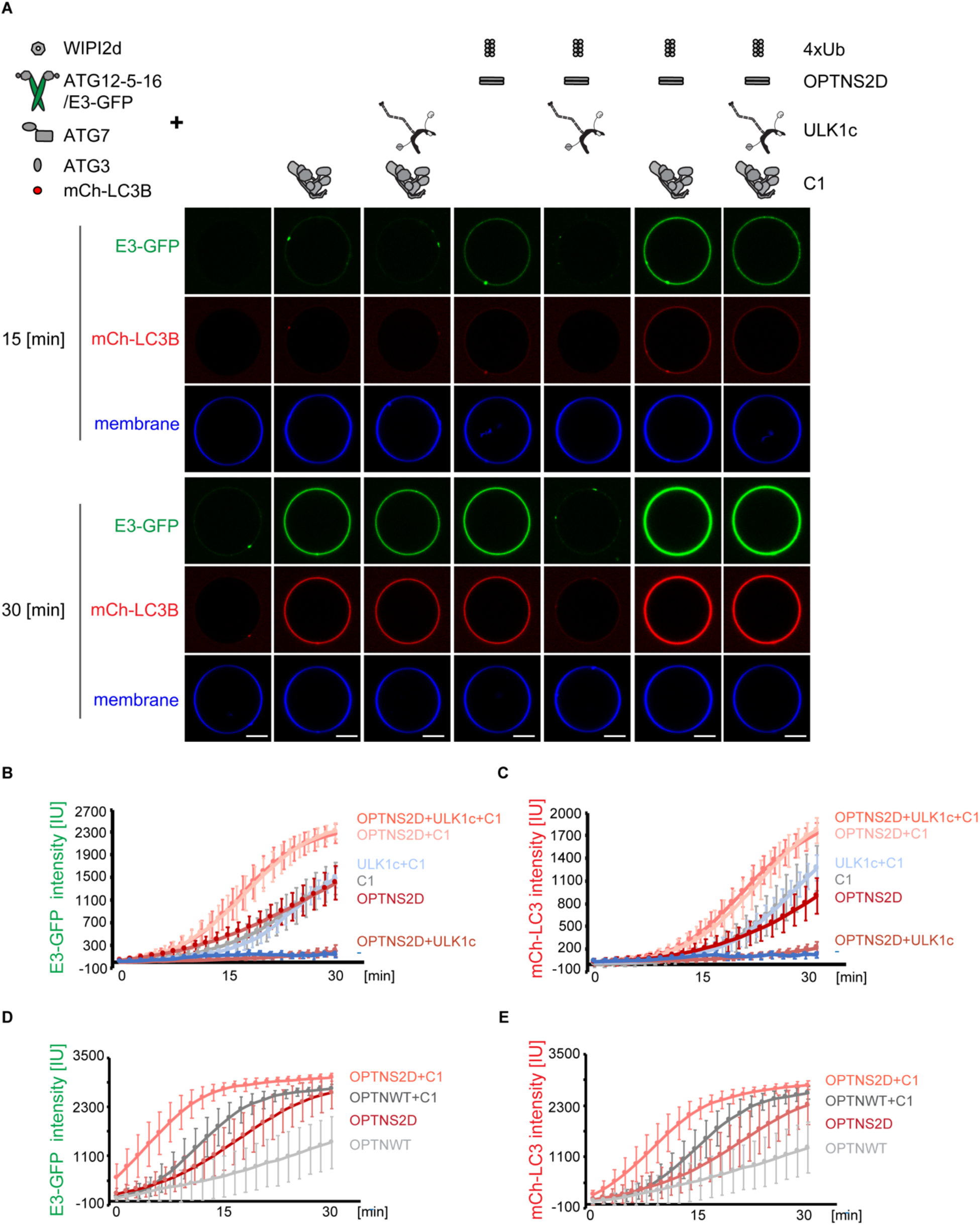
Reconstitution of OPTN triggered LC3 lipidation. (A) Representative confocal images showing the membrane recruitment of E3 complex and LC3B. GUVs were incubated with WIPI2d, E3-GFP, ATG7, ATG3, mCherry-LC3B, ATP/Mn^2+^, and different protein as listed above each image column. Images taken at 15 min and 30 min are shown. Scale bars, 10 μm. (B and C) Quantitation of the kinetics of mCherry-LC3B (B) and E3-GFP (C) recruitment to the membrane from individual GUV tracing in A (Averages of 50 vesicles are shown, error bars indicate standard deviations). (D and E) GUVs were incubated with WIPI2d, E3-GFP, ATG7, ATG3, mCherry-LC3B, GST-Ub_4_, ATP/Mn^2+^, and OPTNWT or OPTNS2D in the presence or absence of PI3KC3-C1. Quantitation of the kinetics of mCherry-LC3B (D) and E3-GFP (E) recruitment to the membrane from individual GUV tracing are shown (Averages of 50 vesicles, error bars indicate standard deviations). All results representative of three independent experiments.

### The kinase activity of ULK1 is dispensable for cargo receptor induced-LC3 lipidation

We next asked whether the kinase activity is required for the receptor induced LC3 lipidation, as the ULK1 kinase has been reported to phosphorylate multiple downstream autophagy components upon autophagy initiation, including the ATG14 and BECN1 subunits of PI3KC3-C1, ATG16L1 and ATG9A (*45, 46, 53–55*). We found that in the presence of NDP52, GST-Ub_4_, PI3KC3-C1, WIPI2d, and the LC3 conjugation machinery, the ULK1 kinase dead (KD) complex accelerated both E3 membrane recruitment and LC3 lipidation to the same extent as the wild-type complex (Fig. 3A and B). In contrast, in the presence of OPTNS2D and all the other components, neither wild-type nor KD ULK1 complex enhanced E3 binding or LC3 lipidation (fig. S5). These data support that OPTN triggered LC3 lipidation is independent of both the catalytic and non-catalytic activities of the ULK1 complex.

**Fig.3.**
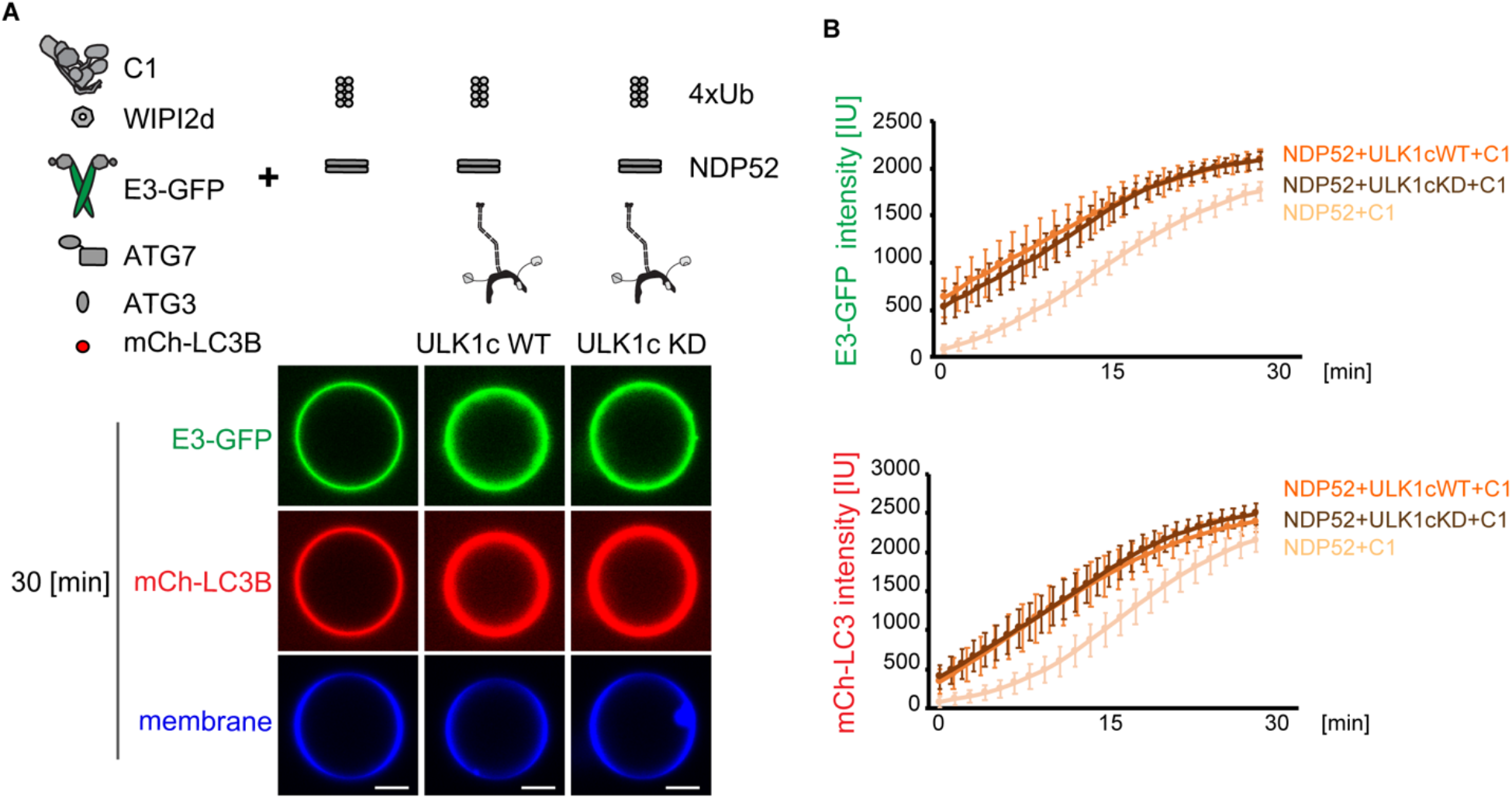
The kinase activity of ULK1 is dispensable for cargo receptor induced-LC3 lipidation. (A) Representative confocal images showing the membrane recruitment of E3-GFP complex and mCherry-LC3B. GUVs were incubated with GST-Ub_4,_ NDP52, PI3KC3-C1, WIPI2d, E3-GFP, ATG7, ATG3, mCherry-LC3B, ATP/Mn^2+^ in the presence or absence of ULK1 WT complex or ULK1 kinase dead complex. Images taken at 30 min are shown. Scale bars, 10 μm. (B) Quantitation of the kinetics of E3-GFP complex and mCherry-LC3B recruitment to the membrane from individual GUV tracing in A are shown (Averages of 50 vesicles, error bars indicate standard deviations). All results representative of three independent experiments.

### Kinetics of ULK1 complex recruitment to membranes

To analyze the differences between the three cargo receptors in more detail, we went on to investigate the kinetics of the recruitment of the upstream components as triggered by cargo receptors. We first monitored the kinetics of ULK1 complex recruitment. In the presence of WIPI2d and conjugation machinery, no detectable GFP tagged ULK1 complex was recruited to membrane (Fig. 4A, first column). The addition of NDP52 or TAX1BP1-Ub, with GST-Ub_4,_ dramatically enhanced the membrane binding of ULK1 complex (Fig. 4A, second and third columns, B, and C), consistent with the previous observations that NDP52 directly recruited ULK1 complex in mitophagy or xenophagy (*28–30*). We noticed that only a little LC3 lipidation occurred even though the ULK1 complex was enriched on the membrane upon the addition of GST-Ub_4_ with either NDP52 or TAX1BP1 (Fig. 4A, second and third columns, B, and C). This suggested that ULK1 complex alone is in-sufficient to activate LC3 conjugation.

**Fig.4.**
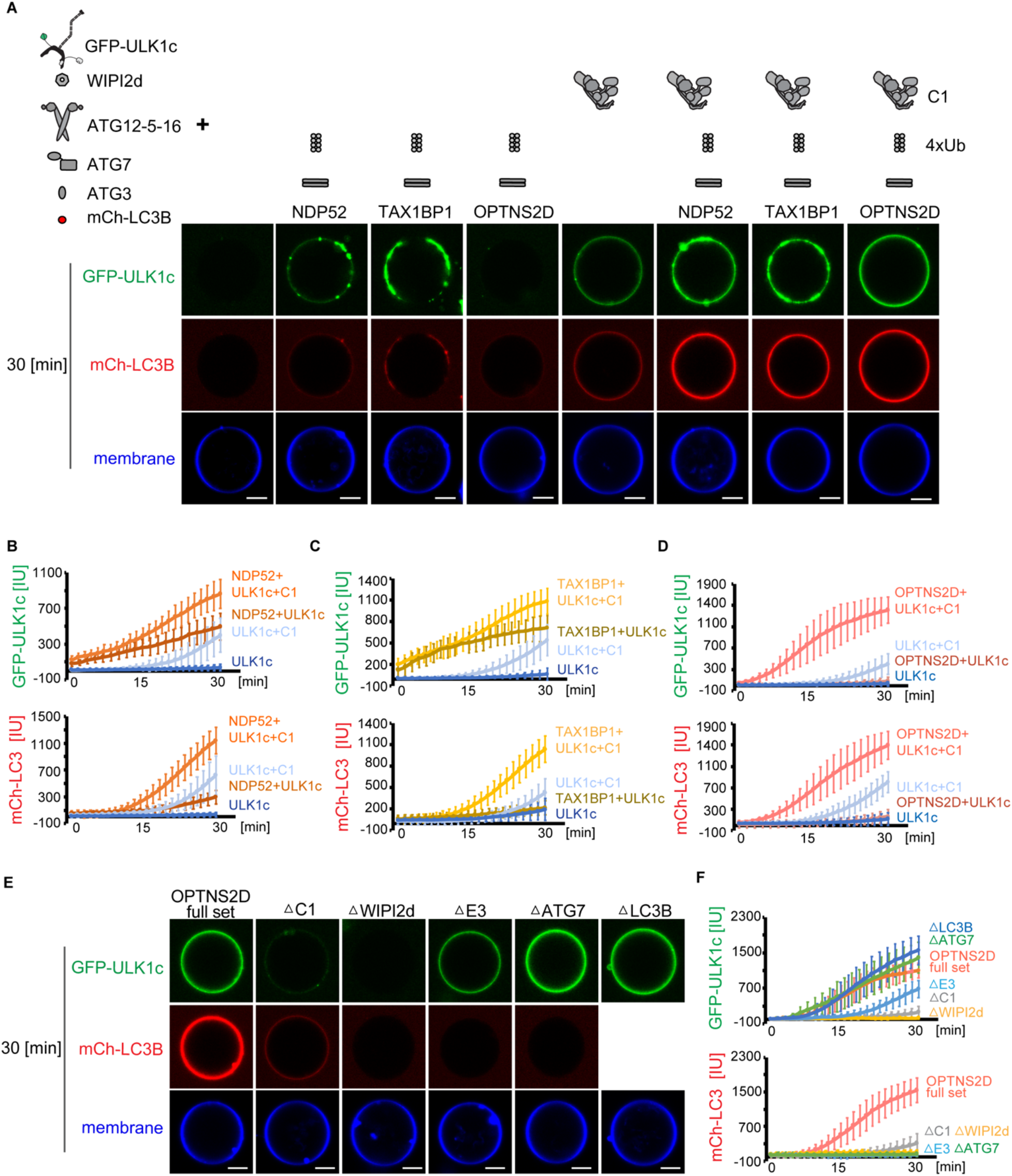
Kinetics of ULK1 complex recruitment to membranes. (A) Representative confocal images showing the membrane recruitment of GFP-ULK1 complex and mCherry-LC3B. GUVs were incubated with WIPI2d, E3 complex, ATG7, ATG3, mCherry-LC3B, ATP/Mn^2+^ and different protein combinations as listed above each image column. Images taken at 30 min are shown. Scale bars, 10 μm. (B-D) GUVs were incubated with WIPI2d, E3 complex, ATG7, ATG3, mCherry-LC3B, ATP/Mn^2+^, and GFP-ULK1 complex or GFP-ULK1 complex together with PI3KC3-C1 complex, in the presence or absence of NDP52 and GST-Ub_4_ (B), or TAX1BP1 and GST-Ub_4_ (C), or OPTNS2D and GST-Ub_4_ (D). Quantitation of the kinetics of GFP-ULK1 complex and mCherry-LC3B recruitment to the membrane from individual GUV tracing are shown (Averages of 50 vesicles, error bars indicate standard deviations). (E) Representative confocal images showing the membrane recruitment of GFP-ULK1 complex and mCherry-LC3B. GUVs were incubated with GST-Ub_4_, OPTNS2D, GFP-ULK1 complex, PI3KC3-C1 complex, WIPI2d, E3 complex, ATG7, ATG3, mCherry-LC3B, and ATP/Mn^2+^ each time omitting one of the components downstream of ULK1 complex. Scale bars, 10 μm. (F) Quantitation of the kinetics of GFP-ULK1 complex and mCherry-LC3B recruitment to the membrane from individual GUV tracing in E are shown (Averages of 50 vesicles, error bars indicate standard deviations). All results representative of three independent experiments.

In contrast to NDP52 or TAX1BP1, the addition of OPTNS2D and GST-Ub_4_ did not result in any increased ULK1 membrane binding (Fig. 4A fourth column, and D). However, when PI3KC3-C1 was added in the reaction, we observed an obvious membrane binding of ULK1 complex, which was further enhanced by the addition of NDP52, TAX1BP1 or even OPTNS2D (Fig. 4A, last four columns, B to D). These data were interpreted in terms of a two-step recruitment of ULK1 complex to membrane in which the ULK1 complex is initially recruited to the membrane by NDP52 or TAX1BP1, but not OPTN. Once PI3KC3-C1 is active on the membrane, ULK1 complex recruitment is promoted further, even when PI3KC3-C1 is recruited downstream of OPTN only.

We then sought to understand the mechanism of PI3KC3-C1 dependent recruitment of the ULK1 as triggered by OPTNS2D. Omission of PI3KC3-C1 or WIPI2d almost completely eliminated the membrane binding of the ULK1 complex. Depletion of E3 complex also largely decreased ULK1 complex recruitment, however, depletion of ATG7 or LC3 slightly increased the ULK1 membrane recruitment (Fig. 4E and F). As expected, omission of any of the components downstream of ULK1 complex eliminated OPTNS2D triggered LC3 lipidation (Fig. 4E and F). These data indicate that the PI3KC3-C1-WIPI2d-E3 axis is required for the further recruitment of ULK1, which is consistent with the previous observations that the translocation of ULK1 complex to omegasomes was stabilized by sustained PI(3)P synthesis (*56*) and that FIP200 could form a trimeric complex with ATG16 and WIPI2 (*57*). The lack of dependence on ATG7 or LC3 rules out that an ULK1 LIR motif-LC3 interaction (*58, 59*) is driving ULK1 recruitment in these experiments. This recruitment of ULK1 complex downstream of OPTN does not lead to a feed forward increase in OPTN triggered LC3 lipidation, given that LC3 lipidation is similar in the presence or absence ULK1 complex (Fig. 2B), which suggests that in OPTN-triggered mitophagy, the ULK1 complex functions at a stage of autophagosome formation subsequent to LC3 lipidation or in other processes that act in parallel to LC3 lipidation.

### Kinetics of PI3KC3-C1 recruitment to membranes

We next monitored the kinetics of PI3KC3-C1 complex recruitment during LC3 lipidation. As distinct from the ULK1 complex, the intrinsic membrane affinity of PI3KC3-C1 enabled it to bind membranes even in the presence of WIPI2d and conjugation machinery but not cargo receptors (Fig. 5A, first column). This is consistent with the observation that PI3KC3-C1 alone can trigger LC3 lipidation in the absence of ULK1 complex (*37*). The addition of OPTNS2D and GST-Ub_4_, but not GST-Ub_4_ with NDP52 or TAX1BP1, dramatically enhanced the membrane binding of PI3KC3-C1 complex (Fig. 5A, first four columns, B to D). ULK1 complex alone did not increase PI3KC3-C1 membrane binding (Fig. 5A, fifth column). However, the addition of ULK1 complex did promote membrane recruitment of PI3KC3-C1 in the presence of NDP52 or TAX1BP1, although not OPTNS2D (Fig. 5A, last three columns, B to D). These data indicate that OPTN strongly enhances membrane recruitment of PI3KC3-C1 on its own. NDP52 and TAX1BP1 have a similar ultimate effect, but only in the presence of ULK1 complex.

**Fig.5.**
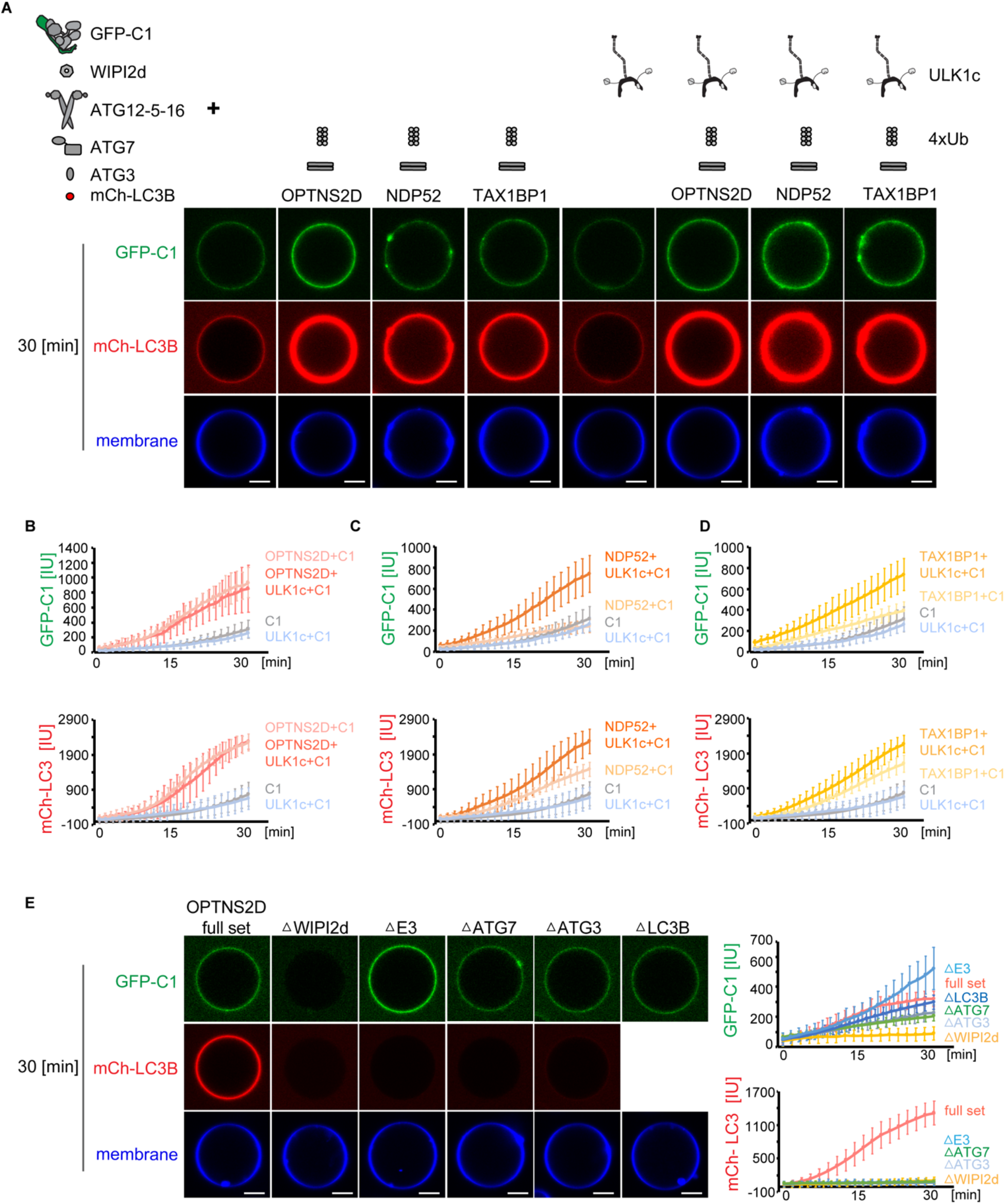
Kinetics of PI3KC3-C1 recruitment to membranes. (A) Representative confocal images showing the membrane recruitment of GFP-PI3KC3-C1 complex and mCherry-LC3B. GUVs were incubated with WIPI2d, E3 complex, ATG7, ATG3, mCherry-LC3B, ATP/Mn^2+^ and different protein combinations as listed above each image column. Images taken at 30 min are shown. Scale bars, 10 μm. (B-D) GUVs were incubated with WIPI2d, E3 complex, ATG7, ATG3, mCherry-LC3B, and GFP-PI3KC3-C1 complex or GFP-PI3KC3-C1 complex together with ULK1 complex, in the presence or absence of OPTNS2D and GST-Ub_4_ (B), or NDP52 and GST-Ub_4_ (C), or TAX1BP1 and GST-Ub_4_ (D). Quantitation of the kinetics of GFP-PI3KC3-C1 complex and mCherry-LC3B recruitment to the membrane from individual GUV tracing are shown (Averages of 50 vesicles, error bars indicate standard deviations). (E) Representative confocal images showing the membrane recruitment of GFP-PI3KC3-C1 complex and mCherry-LC3B. GUVs were incubated with GST-Ub_4_, OPTNS2D, GFP-PI3KC3-C1 complex, WIPI2d, E3 complex, ATG7, ATG3, mCherry-LC3B, and ATP/Mn^2+^ each time omitting one of the components downstream of PI3KC3-C1 complex. Scale bars, 10 μm. (F) Quantitation of the kinetics of GFP-PI3KC3-C1 complex and mCherry-LC3B recruitment to the membrane from individual GUV tracing in E are shown (Averages of 50 vesicles, error bars indicate standard deviations). All results representative of three independent experiments.

Given the dramatic increase of PI3KC3-C1 membrane binding triggered by OPTN, we sought to understand its mechanism. Omission of WIPI2d almost completely blocked the membrane binding of PI3KC3-C1 (Fig. 5E and F), consistent with the previous observation that PI3KC3-C1 and WIPI2d cooperatively bind to membranes (*37*). In contrast, the depletion of ATG7, ATG3, or LC3, but not E3, resulted in a slight decrease of PI3KC3-C1 binding (Fig. 5E and F), suggesting that a multivalent assembly of OPTN, PI3KC3-C1, WIPI2d, and E3 may be responsible for the membrane recruitment of PI3KC3-C1 by OPTN.

### Interactions between cargo receptors and the core autophagy machinery

We found that NDP52, TAX1BP1 and OPTN trigger robust LC3 lipidation by dramatically enhancing the membrane binding of ULK1 or PI3KC3-C1 complex, and that they do so by distinct mechanisms. We therefore hypothesized that these cargo receptors could interact with the autophagy core complexes in distinct ways. To test this, we systematically analyzed the binding between these cargo receptors and different autophagy components by a microscopy-based bead interaction assay (Fig. 6A). The ULK1 complex was specifically recruited to beads coated with NDP52 and TAX1BP1, but not OPTNS2D (Fig. 6B and G), consistent with the observation that NDP52 or TAX1BP1 directly recruited ULK1 complex to membrane. However, no detectable PI3KC3-C1 complex was recruited to beads coated with OPTNS2D. Instead, weak binding between PI3KC3-C1 and NDP52 or TAX1BP1 was detected (Fig. 6C and G), suggesting that the increased membrane binding of PI3KC3-C1 by OPTN was not mediated by a direct interaction. Weak binding between OPTNS2D and WIPI2d, NDP52 and WIPI2d, OPTNS2D and E3, NDP52 and E3 were also observed (Fig. 6D, E and G). We noticed a strong interaction between TAX1BP1 and WIPI2d or E3 (Fig. 6E and G), which may explain the stronger membrane recruitment of E3 by TAX1BP1 and GST-Ub_4_ alone. Interactions between NDP52, TAX1BP1, OPTNS2D and LC3B were weak at the tested concentrations (Fig. 6F and G). These data indicate that cargo receptors directly bind to multiple autophagy components, and thus trigger LC3 lipidation through a multivalent web of both strong and weak interactions.

**Fig. 6.**
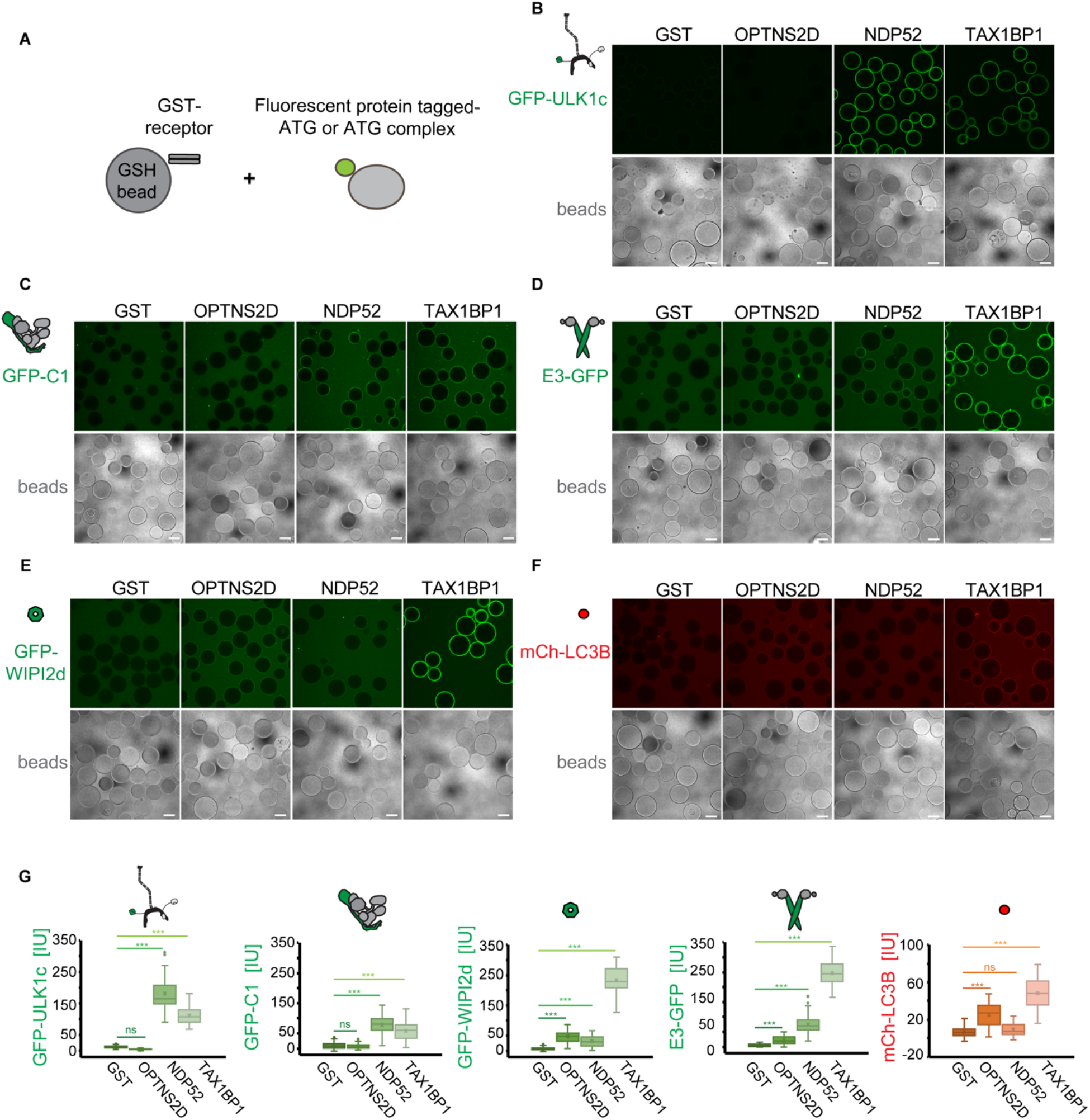
Interactions between cargo receptors and the core autophagy machinery. (A) The schematic drawing illustrates the bead based pull-down setting. (B-F) Representative confocal images showing recruitment of GFP-ULK1 complex (B), GFP-PI3KC3-C1 complex (C), E3-GFP (D), GFP-WIPI2d (E) or mCherry-LC3B (F) to beads coated with GST, GST-NDP52, GST-TAX1BP1 or GST-OPTNS2D. A mixture of GST or GST tagged cargo receptors with different fluorescent protein tagged autophagy components were incubated with GSH beads for 1 h and images were taken and shown. Scale bars, 50 μm. (G) The quantification of GFP or mCherry signal on beads are shown (Averages of 40 beads, error bars indicate standard deviations). All results representative of three independent experiments.

## Discussion

Over the past two years, rapid advances in the mechanistic cell biology of autophagy, and the elucidation of new activities for autophagy proteins, has crystallized into detailed models for mechanisms of autophagosome formation (*4, 60*). The ability to biochemically reconstitute such a pathway from purified components is a stringent and powerful test of such models. Moreover, reconstitution allows nuanced aspects of the interplay between components to be assessed with a rigor that is difficult *in vivo*. In the yeast model system, it was recently shown that a set of purified components could recapitulate the cargo-stimulated Atg8 lipidation and lipid transfer into Atg9 vesicles, confirming the function of Atg9 vesicles as the seeds of the phagophore (*36*). Progress in the reconstitution of human autophagy is less advanced, despite the importance of selective autophagy in many human diseases. We previously reconstituted the PI3KC3-C1, WIPI2, and E3 circuit, demonstrating positive feedback (*37*). Upstream of this circuit, we found that NDP52 mediated the cargo-initiated recruitment of the ULK1 complex to membranes *in vitro* (*29*). Here, we showed that it was possible to reconstitute the circuit connecting the major selective cargo receptors involved in mitophagy, NDP52, TAX1BP1 and OPTN from cargo recognition to LC3 lipidation, with each situation manifesting unique properties (Fig. 7).

**Fig. 7.**
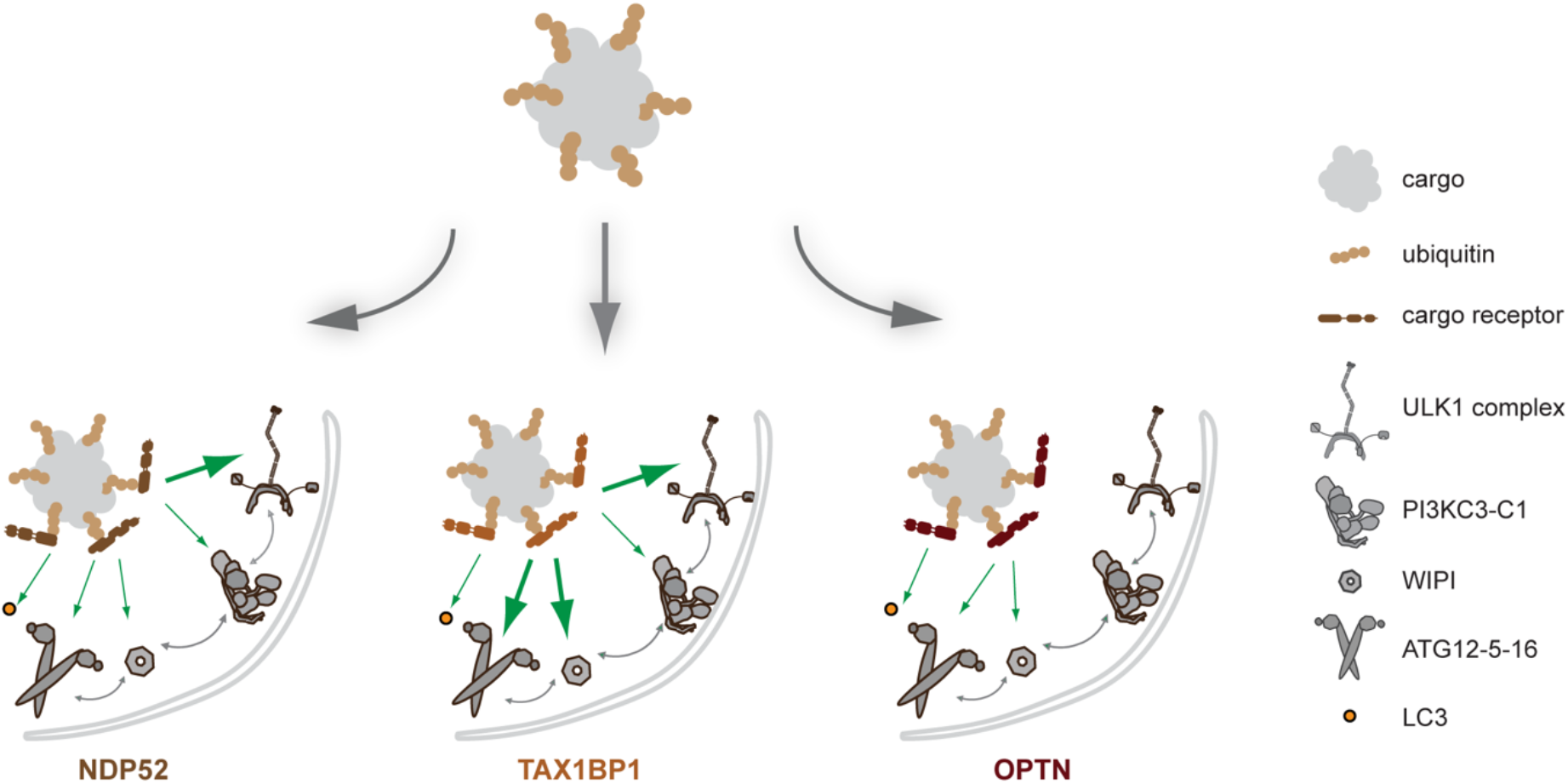
Model for cargo receptor mediated LC3 lipidation. For selective autophagy that degrades targets relying on ubiquitination signals, the cargo receptors like NDP52, TAX1BP1, or OPTN first bind to ubiquitinated cargos, and recruit distinct multiple autophagy machineries through a multivalent web of weak interactions, these components work together to trigger membrane association of LC3 family proteins.

One of the important recent conceptual advances in selective autophagy was the discovery that cargo receptors function upstream of the core autophagy initiating complexes (*28, 30–32*). This paradigm replaced the earlier model that cargo receptors connected substrates to pre-existing LC3-lipidated membranes. Here, we showed that cargo-engaged NDP52, TAX1BP1, or OPTN were capable of potently driving LC3 lipidation in the presence of physiologically plausible nanomolar concentration of the purified autophagy initiation complexes. These reconstitution data directly confirm the new model for cargo-induced formation of LC3-lipidated membranes in human cells.

We found that different cargo receptors use distinct mechanisms to trigger LC3 lipidation downstream of cargo. NDP52 is strongly dependent on the presence of the ULK1 complex, consistent with findings in xenophagy (*28*) and mitophagy (*30*). TAX1BP1 behaves much like NDP52, as expected based on the common presence of an N-terminal SKICH domain, the locus of FIP200 binding (*28*). However, TAX1BP1 was more active in promoting LC3 lipidation in the absence of PI3KC3-C1. Unexpectedly strong binding was observed between TAX1BP1 and E3 and WIPI2d. This raises the possibility that TAX1BP1-mediated selective autophagy may be less dependent on the core complexes as compared to NDP52.

In sharp contrast to NDP52 and TAX1BP1, the *in vitro* LC3 lipidation downstream of OPTN is completely independent of the ULK1 complex. Our finding is consistent with the recent report that OPTN-induced mitophagy, in contrast to NDP52, does not depend on the recruitment of FIP200 (*32*). It is also consistent with our observation of direct binding of NDP52 and TAX1BP1, but not OPTN, to the ULK1 complex *in vitro*. We found that OPTN induced LC3 lipidation *in vitro* is strongly dependent on PI3KC3-C1 and WIPI2d, as expected based on the roles of these proteins in E3 activation. No single one of these complexes, or the E3 itself, bound strongly to OPTN, but all of them bound weakly. This suggests that a multiplicity of weak interactions with several factors contributes to the recruitment of the core complexes downstream of OPTN. ATG9A (*32*), which was not present in this study, likely contributes further to this multivalent web of low affinity interactions.

Subunits of PI3KC3-C1 are phosphorylated by the ULK1 kinase (*45, 55*), and it has long been assumed that these phosphorylation events would promote autophagy. We found, however, that the kinase dead version of the ULK1 complex was as effective in promoting LC3 lipidation as wild-type. This result is consistent with a recent pharmacological study that found ULK1 kinase activity to be dispensable for PI3KC3-C1 activation at p62 condensates (*61*).

In conclusion, we have reconstituted much of the process of cargo-stimulated selective autophagy using purified human proteins. The remaining steps still to be completed *in vitro* are the ATG2 and ATG9-dependent transfer of phospholipids for phagophore growth, and the engulfment of cargo. The observations here provide powerful confirmation for the model that cargo itself triggers formation of LC3-lipidated membranes on a just-in-time basis. They also reveal nuances of how different cargo receptors utilize distinct repertoires of weak and strong interactions with the core complexes to trigger LC3 lipidation. These are subtleties that would have been difficult to uncover in traditional cellular knock out and rescue experiments. These unique modes of core complex recruitment may underlie the divergent core complex phenotypes that are seen in different classes of selective autophagy in different cellular contexts.

## Acknowledgements

This work was supported by the Aligning Science Across Parkinson’s Collaborative Research Network ASAP-0350 (J.H.H. and S.M.), HFSP (RGP0026/2017 to J.H.H. and S.M.), NIH R01 GM111730 (J.H.H.), and the Jane Coffin Childs Foundation (A. L. Y.).

## Materials and Methods

### Plasmid construction

Synthetic codon-optimized DNAs encoding components of human ULK1 complex, PI3KC3-C1 complex were subcloned into the pCAG vector with GST, MBP or TwinStrep-Flag (TSF) tag. Synthetic codon-optimized DNAs encoding human ATG12-ATG5-ATG16 were subcloned into the pGBdest vector with Strep tag. DNA encoding human WIPI2d was subcloned into the pCAG vector with TSF tag. DNA encoding mouse ATG7 was subcloned into the pFast BacHT vector with His tag. DNAs encoding human ATG3, LC3B were subcloned into the pET vector with His tag. DNAs encoding human NDP52, OPTN and TAX1BP1 were subcloned into the pGST2 vector with GST tag. DNA encoding linear tetraubiquitin was subcloned into the pGEX5 vector with GST tag. Details are shown in Table S1.

### Protein expression and purification

The ULK1 complex, PI3KC3-C1 complex and WIPI2d protein were expressed and purified from HEK293 GnTI cells described as previously (*29, 37*). DNAs were transfected cells using polyethylenimine (Polysciences). After 48-72 h expression, cells were harvested and lysed with lysis buffer (50 mM HEPES pH 7.4, 1% Triton X-100, 200 mM NaCl, 1 mM MgCl_2_, 10% glycerol, and 1mM TCEP) supplemented with EDTA free protease inhibitors (Roche). The lysate was clarified by centrifugation (16000 rpm at 4 °C for 1 h) and incubated with resins.

To purify GST-FIP200^Δ641-779^-MBP and GST-FIP200-MBP, the supernatant was incubated with Glutathione Sepharose 4B (GE Healthcare) with gentle shaking at 4 °C for 10 h. The mixture was then loaded onto a gravity flow column, and the resin was washed extensively with wash buffer (50 mM HEPES pH 8.0, 200 mM NaCl, 1 mM MgCl_2_ and 1 mM TCEP). Eluted protein samples flowed through Amylose resin (New England Biolabs) for a second step of affinity purification. The final buffer after MBP affinity purification is 20 mM HEPES pH 8.0, 200 mM NaCl, 2 mM MgCl_2_, 1 mM TCEP and 50 mM maltose. To purify (±GFP)-ULK1 complex for studying the effect of FIP200^Δ641-779^ and ULK1 (kinase-dead mutant), FIP200/ATG13/ATG101 subcomplex and ULK1 were expressed and purified separately. After the first step of affinity purification, the two samples were mixed, cleaved by TEV at 4 °C overnight, and subjected to a second step of affinity purification using the MBP tag. The final buffer after MBP affinity purification is 20 mM HEPES pH 8.0, 200 mM NaCl, 2 mM MgCl_2_, 1 mM TCEP and 50 mM maltose. For the rest of GUV experiments, FIP200/ATG13/ATG101 subcomplex and ULK1 were expressed and purified separately in both first and second steps of affinity purification. The final buffer after second step MBP affinity purification is 20 mM HEPES pH 8.0, 200 mM NaCl, 2 mM MgCl_2_, 1 mM TCEP and 50 mM maltose. The complexes were used immediately for the GUV assays.

To purify (±GFP)-PI3KC3-C1 complex, the supernatant was incubated with Glutathione Sepharose 4B (GE Healthcare) at 4 °C for 4 h, applied to a gravity column, and washed extensively with wash buffer (50 mM HEPES pH 8.0, 200 mM NaCl, 1 mM MgCl_2_, and 1mM TCEP). The protein complexes were eluted with wash buffer containing 50 mM reduced glutathione, and then treated with TEV protease at 4 °C overnight. TEV-treated complexes were loaded on a Strep-Tactin Sepharose gravity flow column (IBA, GmbH). The complexes were eluted with a final buffer containing 20 mM HEPES pH 8.0, 200 mM NaCl, 2 mM MgCl_2_, 1 mM TCEP, and 10 mM desthiobiotin (Sigma), and then used immediately for the GUV assays.

To purify (±GFP)-WIPI2d protein, the supernatant was incubated with Strep-Tactin Sepharose resin at 4 °C for 3 h, applied to a gravity column, and washed extensively with wash buffer (50 mM HEPES pH 7.5, 200 mM NaCl, and 1mM TCEP). The proteins were eluted with wash buffer containing 10 mM desthiobiotin, applied onto a Superdex 200 column (16/60 prep grade, GE Healthcare). The final buffer after gel filtration is 20 mM HEPES pH 7.5, 200 mM NaCl, and 1 mM TCEP. Fractions containing pure (±GFP)-WIPI2d protein were pooled, concentrated, snap frozen in liquid nitrogen and stored at −80 °C.

The ATG12-ATG5-ATG16 complex and ATG7 protein were expressed and purified from Sf9 cells as previously described (*37*). Sf9 cells were infected with a single virus stock P1 corresponding to the poli-cystronic construct coding ATG12-ATG5-ATG16 complex or ATG7. Cells were harvested 72 h after infection, lysed and clarified following the same procedure for mammalian cells described as above. To purify (±GFP)-ATG12-ATG5-ATG16 complex, the supernatant was incubated with Strep-Tactin Sepharose at 4 °C for 3 h, applied to a gravity column, and washed extensively with wash buffer (50 mM HEPES pH 7.5, 200 mM NaCl, and 1mM TCEP).

The proteins were eluted with wash buffer containing 10 mM desthiobiotin, applied onto a Superdex 6 column (10/300 Increase). The final buffer after gel filtration is 20 mM HEPES pH 7.5, 200 mM NaCl, and 1 mM TCEP. Peak fractions containing pure (±GFP)-ATG12-ATG5-ATG16 complexes were pooled and snap frozen in liquid nitrogen and stored at −80 °C.

To purify ATG7 protein, the supernatant was loaded on a Ni-NTA column (GE Healthcare) gravity flow column, washed extensively with wash buffer (50 mM HEPES pH 7.5, 200 mM NaCl, 20 mM imidazole and 1mM TCEP). The proteins were eluted with wash buffer containing 200 mM imidazole, applied onto a Superdex 200 column (16/60 prep grade). The final buffer after gel filtration is 20 mM HEPES pH 7.5, 200 mM NaCl, and 1 mM TCEP. Peak fractions containing pure ATG7 protein were pooled and snap frozen in liquid nitrogen and stored at −80 °C.

The linear tetraubiquitin, NDP52, OPTN, TAX1BP1, ATG3 and mCherry-LC3B were expressed and purified from *E. coli* (BL21DE3). Protein expression was induced with 100 μM IPTG when cells were grown to an OD600 of 0.8 and further grown at 18°C overnight. Cells were harvested and stocked in −80 °C if needed. To purify GST tagged linear tetraubiquitin and receptors, the pellets were resuspended in a buffer containing 50 mM HEPES pH 7.5, 300 mM NaCl, 1 mM TCEP and protease inhibitors (Roche), and sonicated before being cleared at 16000 rpm at 4 °C for 1 h. The supernatant was incubated with Glutathione Sepharose 4B at 4 °C for 4 h, applied to a gravity column, and washed extensively with wash buffer (50 mM HEPES pH 7.5, 300 mM NaCl, and 1mM TCEP). The proteins were eluted with wash buffer containing 50 mM reduced glutathione, and then applied onto a Superdex 6 column (10/300 Increase). The final buffer after gel filtration is 20 mM HEPES pH 8.0, 200 mM NaCl, and 1 mM TCEP. Peak fractions containing pure proteins were pooled and snap frozen in liquid nitrogen and stored at −80 °C.

To purify ATG3 and mCherry-LC3B, the pellets were resuspended in a buffer containing 50 mM HEPES pH 7.5, 300 mM NaCl, 1 mM TCEP, 20 mM imidazole and protease inhibitors, sonicated and clarified. The supernatant was loaded on a Ni-NTA column (GE Healthcare) gravity flow column, washed extensively with wash buffer (50 mM HEPES pH 7.5, 300 mM NaCl, 20 mM imidazole and 1mM TCEP). The proteins were eluted with wash buffer containing 200 mM imidazole, applied onto a Superdex 75 column (16/60 prep grade). The final buffer after gel filtration is 20 mM HEPES pH 7.5, 200 mM NaCl, and 1 mM TCEP. Fractions containing pure proteins were pooled, concentrated, snap frozen in liquid nitrogen and stored at −80 °C.

### Preparation of Giant Unilamellar Vesicles (GUVs)

GUVs were prepared by hydrogel-assisted swelling as described previously (*37*). Briefly, 60 μL 5% (w/w) polyvinyl alcohol (PVA) with a molecular weight of 145,000 (Millipore) was coated onto a plasma-cleaned coverslip of 25 mm diameter. The coated coverslip was placed in a heating incubator at 60 °C to dry the PVA film for 30 min. For all the GUV experiments, a lipid mixture with a molar composition of 64.8% DOPC, 20% DOPE, 10% POPI, 5% DOPS and 0.2% Atto647N DOPE at 1 mg/ml was spread uniformly onto the PVA film. The lipid-coated coverslip was then put under vacuum overnight to evaporate the solvent. 400 μL 400 mOsm sucrose solution was used for swelling for 1 h at room temperature, and the vesicles were then harvested and used within 12 h.

Atto647N DOPE (Atto TEC) was used as the GUV membrane dye. All the other lipids for GUVs preparation are from Avanti Polar Lipids.

### In vitro reconstitution GUV assay

The reactions were set up in an eight-well observation chamber (Lab Tek) at room temperature. The chamber was coated with 5 mg/ml β casein for 30 min and washed three times with reaction buffer (20 mM HEPES at pH 8.0, 190 mM NaCl and 1 mM TCEP). A final concentration of 5 μM GST-4xUb, 500 nM cargo receptors, 25 nM ULK1 complex, 25 nM PI3KC3-C1 complex, 100 nM WIPI2d, 50 nM ATG12-ATG5-ATG16 complex, 100 nM ATG7, 100 nM ATG3, 500 nM mCherry-LC3B, 50 μM ATP, and 2 mM MnCl_2_ was used for all reactions unless otherwise specified. 10 μL GUVs were added to initiate the reaction in a final volume of 120 μL. After 5 min incubation, during which random views were picked for imaging, time-lapse images were acquired in multitracking mode on a Nikon A1 confocal microscope with a 63 × Plan Apochromat 1.4 NA objective. Three biological replicates were performed for each experimental condition. Identical laser power and gain settings were used during the course of all conditions.

### Microscopy-based bead protein-protein interaction assay

A mixture of 0.5 μM GST or GST tagged cargo receptors and different ATG proteins was incubated with 10 μL Glutathione Sepharose beads (GE Healthcare) in a reaction buffer containing 20 mM HEPES at pH 8.0, 200 mM NaCl and 1 mM TCEP. The final concentration of different ATG proteins was as following: 25 nM GFP-ULK1 complex, 25 nM GFP-PI3KC3-C1 complex, 100 nM GFP-WIPI2d, 50 nM ATG12-ATG5-ATG16-GFP complex, and 500 nM mCherry-LC3B.

After incubation at room temperature for 60 min, the beads were washed three times, suspended in 120 μL reaction buffer, and then transferred to the observation chamber for imaging. Images were acquired on a Nikon A1 confocal microscope with a 63 × Plan Apochromat 1.4 NA objective. Three biological replicates were performed for each experimental condition.

### Negative Stain Electron Microscopy Preparation, Collection and Coiled-coil Tracing

Protein sample of GST-FIP200^Δ641-779^-MBP was diluted to 100 nM concentration in elution buffer. 5 μL of sample was applied to continuous carbon grids which were glow discharge in a PELCO easiGlow instrument for 45 s at 25 mAmps. Protein was wicked away using torn 597 Whatman paper and immediately stained with 2% Uranyl acetate. Wicking was repeated again for a second round of 2% uranyl acetate staining.

Data was collected at 120 kV on a Tecnai T12 microscope with a nominal magnification of 49000x. 25 micrographs were taken with a Gatan CCd 4k × 4k camera at a pixel size of 2.2 Å/pixel. Protein particles were manually selected using the manual picking tool within Relion 3.1 and extracted at a binned box size of 120 by 120 corresponding to a pixel size of 8.8 Å/pixel. Extracted particles were measured for coli-coli length in FIJI as previously described (*29*). In brief, 96 single particles were traced using the Simple Neurite Tracer plug in for FIJI. Histogram of the data was prepared for both path length of the coli-coli (82 nm) and the end to end distance of the coli-coli (62 nm).

### Image quantification

GUV images were analyzed using a custom script implemented in Python 3.6 (https://github.com/Hurley-Lab/GUVquantification/blob/main/GUVintensity-2channel.ipynb). Briefly, to obtain the outline of all the vesicles within a field of view, images were segmented into regions corresponding to local maxima of the membrane fluorescence channel, which were defined by applying an Otsu threshold to the differences between local maxima and minima. Then, binding of the fluorescently labelled proteins was quantified by taking the mean value of these segmented pixels in the fluorescent protein channel. Background was calculated as the average of the vesicle-internal background and the vesicle-external background and subtracted from the fluorescence signal. The intensity trajectories of multiple fields of view were then obtained frame by frame. Multiple intensity trajectories were calculated, and the averages and standard deviations were calculated and reported.

For quantification of protein intensity binding to bead, the outline of individual bead was manually defined based on the bright field channel. The intensity threshold was calculated by the average intensities of pixels inside and outside of the bead and then intensity measurements of individual bead were obtained. Averages and standard deviations were calculated among the measured values per each condition and plotted in a bar graph.

## Statistical analysis

Statistical analysis was performed by unpaired Student’s t test using GraphPad Prism 9. P < 0.05 was considered statistically significant.

## Supplementary Materials

**Fig. S1.**
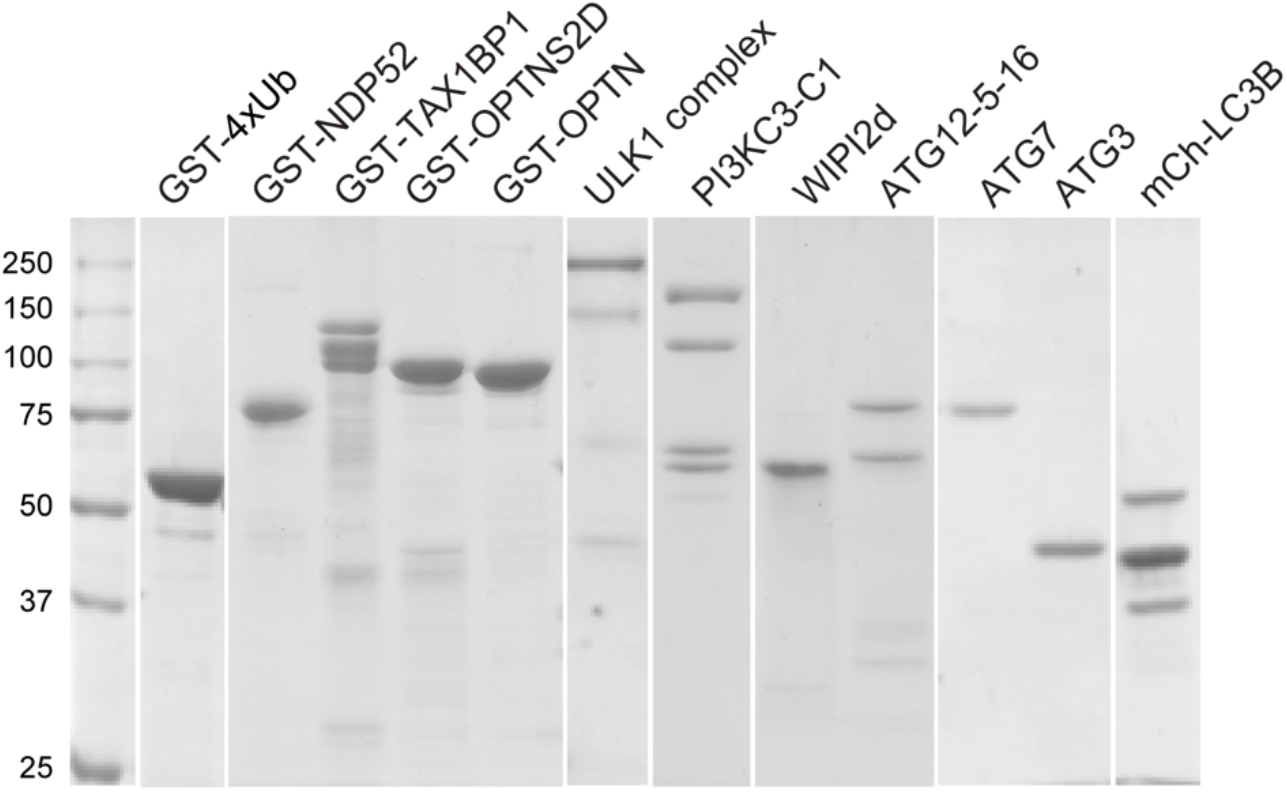
Purification of core autophagy machinery. All purified autophagy components were resolved on a 10% SDS PAGE and shown by Coomassie Brilliant Blue stain.

**Fig. S2.**
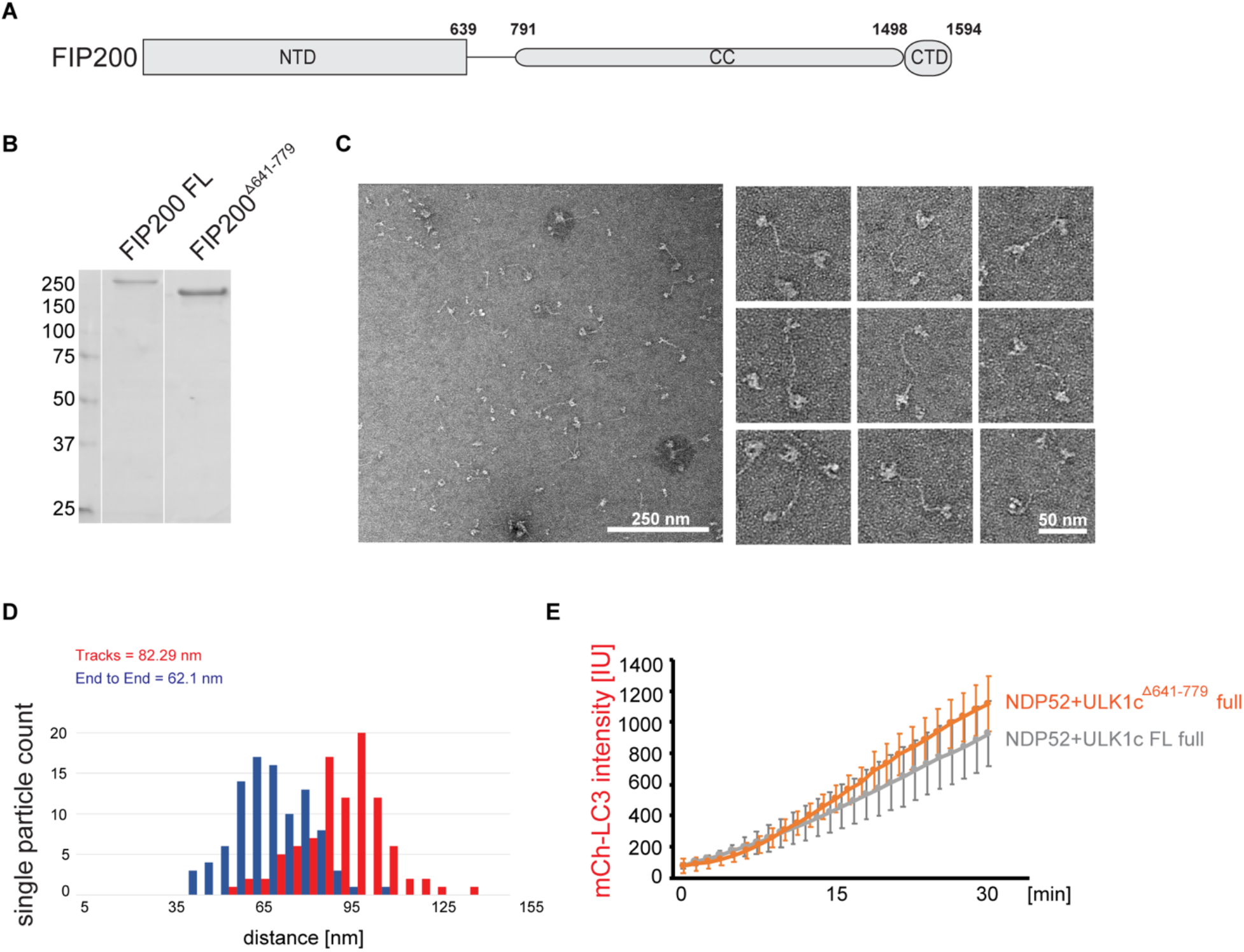
Characterization of ULK1 complex with FIP200^Δ641-779^. (A) The schematic drawing shows the domain structure of FIP200. (B) Purified FIP200 full-length or FIP200^Δ641-779^ was resolved on a 10% SDS PAGE and shown by Coomassie Brilliant Blue stain. (C) Negative stain EM single particles of FIP200^Δ641-779^. (D) Histogram of FIP200^Δ641-779^ path length and end-to-end distances. (E) GUVs were incubated with GST-Ub_4_, NDP52, PI3KC3-C1, WIPI2d, E3, ATG7, ATG3, mCherry-LC3B, ATP/Mn^2+^ in the presence of ULK1 WT complex or ULK1 complex with FIP200^Δ641-779^. Quantitation of the kinetics of mCherry-LC3B recruitment to the membrane from individual GUV tracing are shown (Averages of 50 vesicles, error bars indicate standard deviations).

**Fig.S3.**
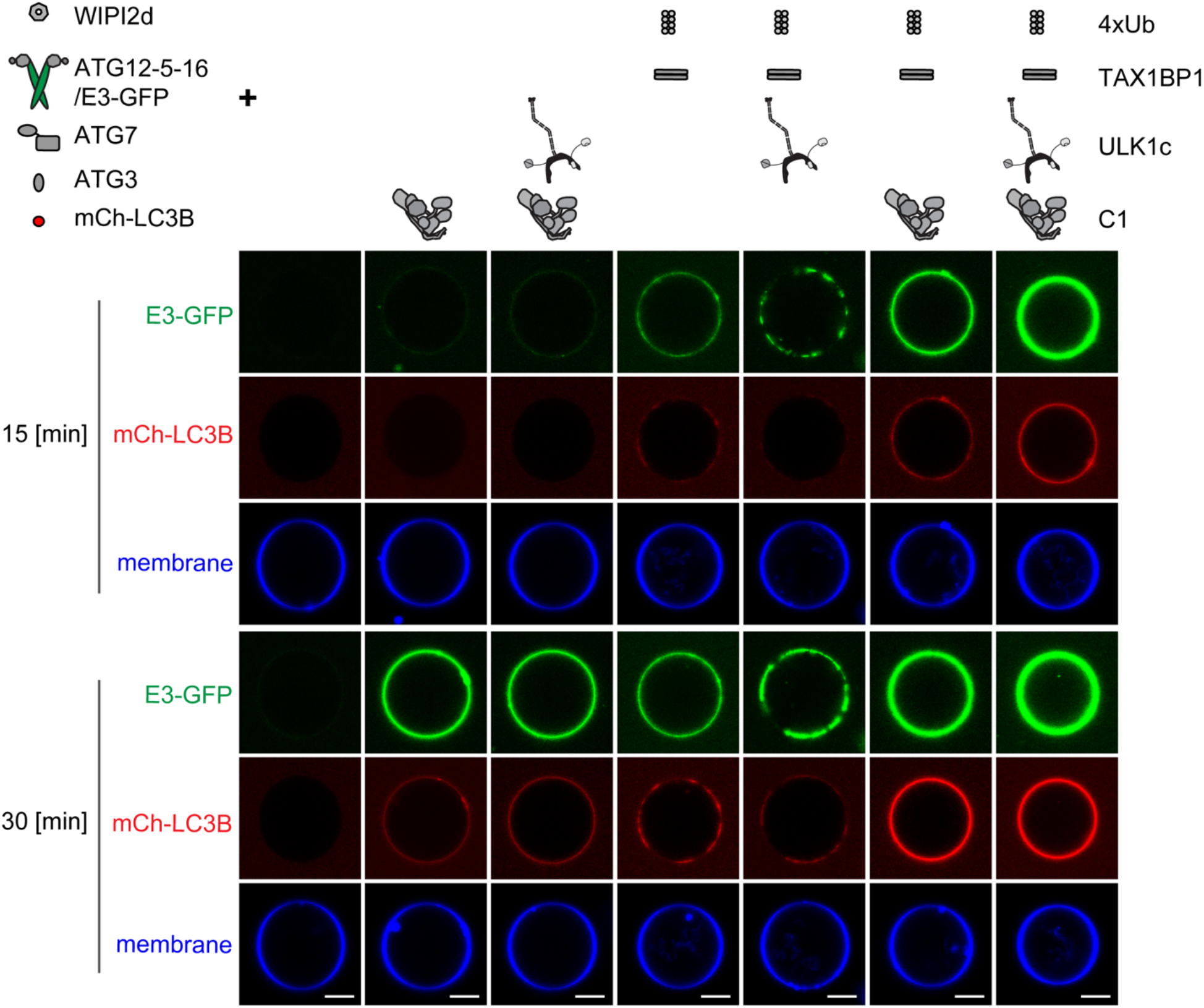
Reconstitution of TAX1BP1-triggered LC3 lipidation. Representative confocal images showing the membrane recruitment of E3-GFP and mCherry-LC3B. GUVs were incubated with WIPI2d, E3-GFP, ATG7, ATG3, mCherry-LC3B, ATP/Mn^2+^, and different upstream components as listed above each image column, respectively. Images taken at 15 min and 30 min are shown. Scale bars, 10 μm.

**Fig. S4.**
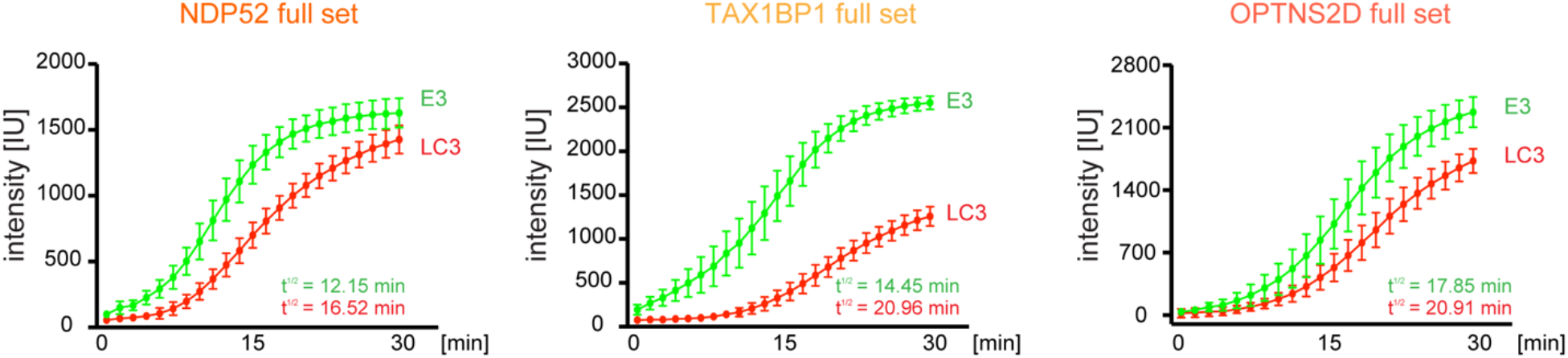
The kinetics of E3 and LC3 membrane recruitment. Quantitation of the kinetics of E3-GFP and mCherry-LC3B recruitment to the membrane from individual GUV tracing are shown (Averages of 50 vesicles, error bars indicate standard deviations). The GUVs were incubated with GST-Ub_4_, ULK1 complex, PI3KC3-C1, WIPI2d, E3, ATG7, ATG3, mCherry-LC3B, ATP/Mn^2+^ in the presence of NDP52, TAX1BP1 or OPTNS2D. The data were fitted into the Boltzmann sigmoidal curve by GraphPad Prism 9, and t^1/2^ was calculated.

**Fig. S5.**
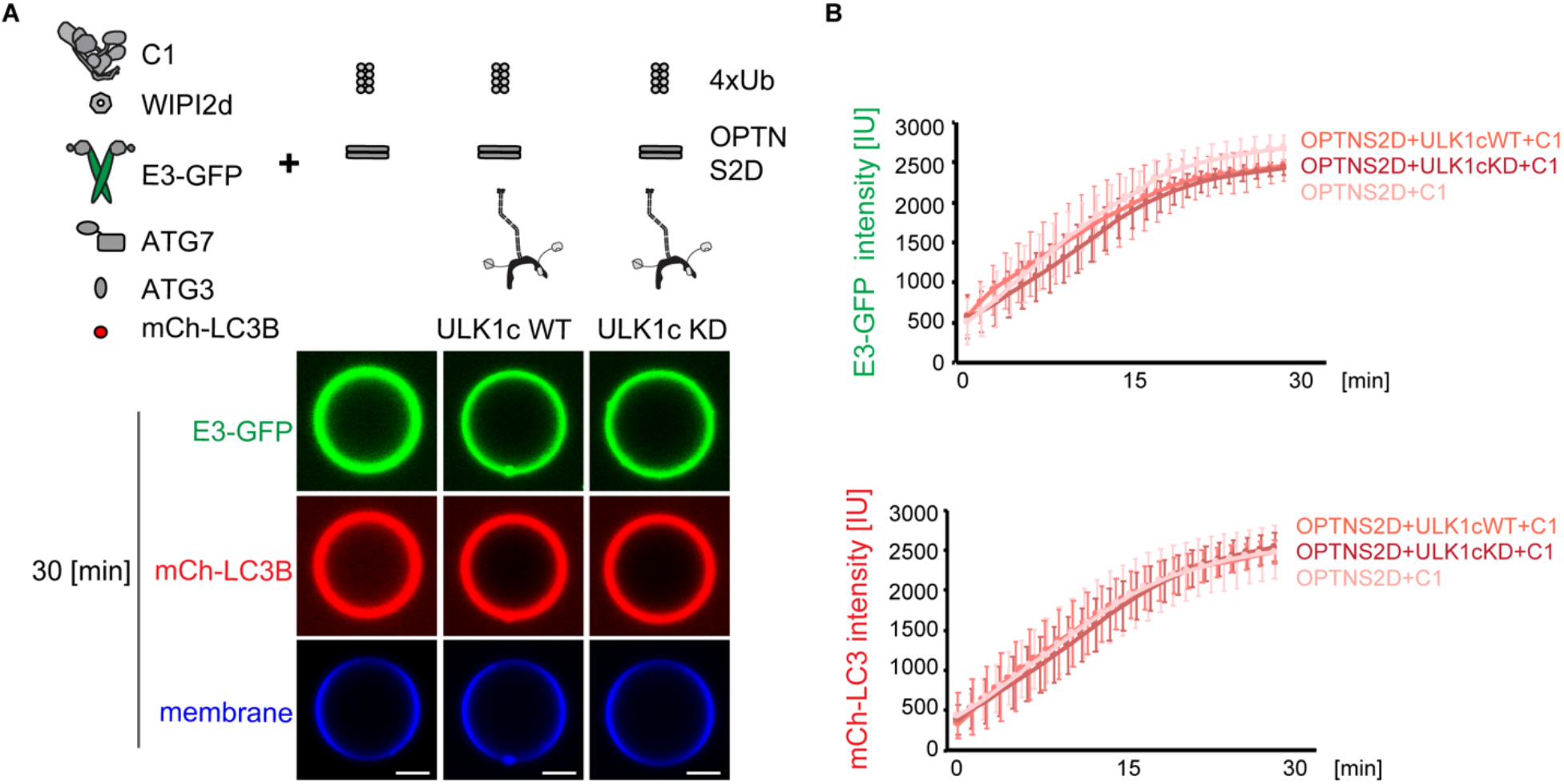
The kinase activity of ULK1 is dispensable for OPTN induced-LC3 lipidation. (A) Representative confocal images showing the membrane recruitment of E3-GFP complex and mCherry-LC3B. GUVs were incubated with GST-Ub_4,_ OPTNS2D, PI3KC3-C1, WIPI2d, E3-GFP, ATG7, ATG3, mCherry-LC3B, ATP/Mn^2+^ in the presence or absence of ULK1 WT complex or ULK1 kinase dead complex. Images taken at 30 min are shown. Scale bars, 10 μm. (B) Quantitation of the kinetics of E3-GFP complex and mCherry-LC3B recruitment to the membrane from individual GUV tracing in C are shown (Averages of 50 vesicles, error bars indicate standard deviations). All results representative of three independent experiments.

**Table S1.**

| **Construct** | **Vector** | **Expression system** | **Published** |
| --- | --- | --- | --- |
| GST-TEVcs-FIP200-MBP | pCAG | HEKGnTi | (29) |
| GST-TEVcs- FIP200<sup>Δ641-779</sup>-MBP | pCAG | HEKGnTi | This study |
| ATG13 | pCAG | HEKGnTi | (29) |
| GFP-ATG13 | pCAG | HEKGnTi | (29) |
| GST- TEVcs-ATG101 | pCAG | HEKGnTi | (29) |
| GST-TEVcs-GFP-ATG101 | pCAG | HEKGnTi | (29) |
| MBP-TSF-TEVcs-ULK1 | pCAG | HEKGnTi | (29) |
| MBP-TSF-TEVcs-ULK1K46I | pCAG | HEKGnTi | This study |
| GST-TEVcs-ATG14 | pCAG | HEKGnTi | (62) |
| GST-TEVcs-GFP-ATG14 | pCAG | HEKGnTi | (37) |
| TSF-TEVcs-VPS34 | pCAG | HEKGnTi | (37) |
| TSF-TEVcs-BECN1 | pCAG | HEKGnTi | (62) |
| VPS15 | pCAG | HEKGnTi | (63) |
| WIPI2d-TEVcs-TSF | pCAG | HEKGnTi | (37) |
| GFP-WIPI2d-TEVcs-TSF | pCAG | HEKGnTi | This study |
| ATG12-10xHis-TEVcs-ATG5-10xHis-TEVcs-ATG16L1-TEVcs-StrepII-ATG7-ATG10 | pGBdest | Sf9 | (37) |
| ATG12-10xHis-TEVcs-ATG5-10xHis-TEVcs-ATG16L1-GFP-TEVcs-StrepII-ATG7-ATG10 | pGBdest | Sf9 | (37) |
| 6xHis-TEVcs-ATG7 | pFast<br>BacHT(B) | Sf9 | (37) |
| 6xHis-TEVcs-ATG3 | pET Duet-1 | E. coli | (37) |
| 6xHis-TEVcs-mCherry-LC3B-Gly(Δ5C) | pET Duet-1 | E. coli | (37) |
| GST-Ub<sub>4</sub> | pGEX5 | E. coli | (29) |
| GST-NDP52 | pGST2 | E. coli | (29) |
| GST-TAX1BP1 | pGST2 | E. coli | This study |
| GST-OPTN | pGST2 | E. coli | This study |
| GST-OPTNS177DS473D | pGST2 | E. coli | This study |

